# Discovery of synthetic lethal and tumor suppressive paralog pairs in the human genome

**DOI:** 10.1101/2020.12.20.423710

**Authors:** Phoebe C. R. Parrish, James D. Thomas, Shriya Kamlapurkar, Austin Gabel, Robert K. Bradley, Alice H. Berger

**Author notes:** Corresponding author: Alice Berger.

## Abstract

CRISPR knockout screens have accelerated the discovery of important cancer genetic dependencies. However, traditional CRISPR-Cas9 screens are limited in their ability to assay the function of redundant or duplicated genes. Paralogs in multi-gene families constitute two-thirds of the protein-coding genome, so this blind spot is the rule, not the exception. To overcome the limitations of single gene CRISPR knockout screens, we developed <u>p</u>aired <u>g</u>uide RNAs for <u>P</u>aralog g<u>EN</u>etic interaction mapping (pgPEN), a pooled CRISPR/Cas9 approach which targets over a thousand duplicated human paralogs in single knockout and double knockout configurations. We applied pgPEN to two cell lineages and discovered that over 10% of human paralogs exhibit synthetic lethality in at least one cellular context. We recovered known synthetic lethal paralogs such as *MAP2K1/MAP2K2*, important drug targets such as *CDK4/CDK6*, and numerous other synthetic lethal pairs such as *CCNL1/CCNL2.* In addition, we identified ten tumor suppressive paralog pairs whose compound loss promotes cell growth. These findings identify a large number of previously unidentified essential gene families and nominate new druggable targets for oncology drug discovery.

**Highlights:**

- Comprehensive genetic interaction mapping of 1,030 human duplicated paralogs using a dual targeting CRISPR/Cas9 approach
- Duplicated paralogs are highly enriched for genetic interactions
- Synthetic lethal paralogs include *CCNL1/CCNL2, CDK4/CDK6*, and *GSK3A/GSK3B*
- Tumor suppressor paralog pairs include *CDKN2A/CDKN2B* and *FBXO25/FBXO32*

## Introduction

CRISPR-Cas9 technology has revolutionized functional genomics by enabling high-fidelity, genome-scale, multiplexed loss-of-function screens in human cells. Due to high specificity and ease of application, genome-wide CRISPR screens are increasingly used to identify cancer drug targets and determine mechanisms of drug resistance (Bartha et al., 2018; Blomen et al., 2015; Hart et al., 2015; Tsherniak et al., 2017; Wang et al., 2017, 2015). However, single-gene knockout studies have a major blind spot: they are unable to assay the function of paralogs — ancestrally duplicated genes that frequently retain functional redundancy. The human genome exhibits a high degree of redundancy as a result of diploidy, gene duplication, and functional overlap of metabolic and signaling pathways (Dean et al., 2008; Harrison et al., 2007; Lavi, 2015; Ohno, 2013). Remarkably, paralogs constitute two-thirds of the human genome, making this blind spot the rule, not an exception, and paralogous genes are less likely to be essential for cell growth than non-paralogous (“singleton”) genes in CRISPR knockout screens (Wang et al., 2015). This blind spot therefore obscures our understanding of normal human genome function and impedes the identification of new cancer drug targets.

Genetic interactions between paralogs have been extensively characterized in yeast, revealing fundamental insights about the differences between whole-genome and small-scale duplicates, functional groups that are enriched for interacting paralogs, and paralog mRNA expression patterns (Dean et al., 2008; Diss et al., 2017; Guan et al., 2007; Harrison et al., 2007). Essential paralogs that compensate for one another’s function exhibit “synthetic lethality,” a genetic interaction in which elimination of the entire family is deleterious but individual loss is tolerated. Yeast geneticists have defined quantitative measures of genetic interactions, which can capture both positive (buffering) and negative (synthetic lethal) interactions (Collins et al., 2006). While paralog genetic interactions are still poorly characterized in mammalian cells, the extensive degree of duplication in the human genome is similar to that seen in yeast (Dennis and Eichler, 2016; Lan and Pritchard, 2016; Singh et al., 2012), so experimental evaluation of human cells is likely to also reveal complex genetic interactions.

Querying the genetic interaction space of the human genome has been limited by current technology; to survey even every possible pairwise interaction, let alone higher order interactions, would involve ~400 million unique genetic perturbations. Moreover, the landscape of genetic interaction among randomly selected genes is exceedingly sparse; existing studies of much smaller sets of gene pairs in human cells identified genetic interactions in less than 0.1% of unrelated gene pairs (Han et al., 2017). To proactively identify these rare but functionally important interactions, research should therefore focus on high-value sets of genes likely to be enriched for functional interactions, such as paralogs.

Interestingly, the same duplication that makes paralogs difficult to study provides a tactical advantage for cancer therapy: the highly rearranged genomes typical of cancer often harbor paralog deletions and inactivating mutations. Cancer-associated loss-of-function of one paralog can confer a dependency on the continued activity of a duplicated pair (De Kegel and Ryan, 2019; Lord et al., 2020; Viswanathan et al., 2018) and this phenomenon has been used to identify synthetic lethal relationships of paralogs such as *MAGOH/MAGOHB* (Viswanathan et al., 2018), *ARID1A/ARID1B* (Helming et al., 2014), and *SMARCA2/SMARCA4* (Hoffman et al., 2014). If the remaining actively expressed paralog could be targeted in tumors with loss of its pair, then tumor cells may show a selective therapeutic window compared to the surrounding normal cells with expression of both paralog members. A successful example of a therapy based on a synthetic lethal interaction is the enhanced sensitivity to PARP inhibitors in *BRCA1-* and *BRCA2-*mutant tumors (Bryant et al., 2005; Farmer et al., 2005; Lord and Ashworth, 2017). We hypothesized that paralogs could provide a rich source of genetic interactions and that direct experimental identification of synthetic lethal paralogs could therefore enable future drug discovery efforts.

Recently, several groups have developed innovative methods for assessing human genetic interactions (GIs) at scale (Boettcher et al., 2018; Dede et al., 2020; DeWeirdt et al., 2020; Gier et al., 2020; Gonatopoulos-Pournatzis et al., 2020; Han et al., 2017; Horlbeck et al., 2018; Najm et al., 2018; Shen et al., 2017). Consistently, while the overall rate of genetic interaction among gene pairs is low, many of the GIs identified were in paralogous genes. To comprehensively identify GIs between human paralogs, we here report our direct experimental evaluation of genetic interactions among 1,030 paralog pairs (2,060 genes) in two human cell contexts. Our analysis revealed not only an extraordinarily high rate of paralog synthetic lethality, but also identified positive interactions that nominate ten paralog pairs as tumor suppressor gene families.

## Results

### A paralog blind spot limits discovery of essential genes and cancer dependencies

The human genome is highly duplicated, with paralogous genes constituting over two thirds of protein coding genes (**Fig. 1A**). Like other groups (Dandage and Landry, 2019; Dede et al., 2020; Wang et al., 2015), we noticed that paralogous genes are less likely to be essential for cell growth than non-paralogous “singleton” genes in CRISPR knockout screening data (**Fig. 1B**) (Vichas et al., 2020). Given the utility of targeting cancer-essential genes for therapy, we reasoned that this paralog blind spot may prevent detection of important druggable cancer dependencies.

**Figure 1.**
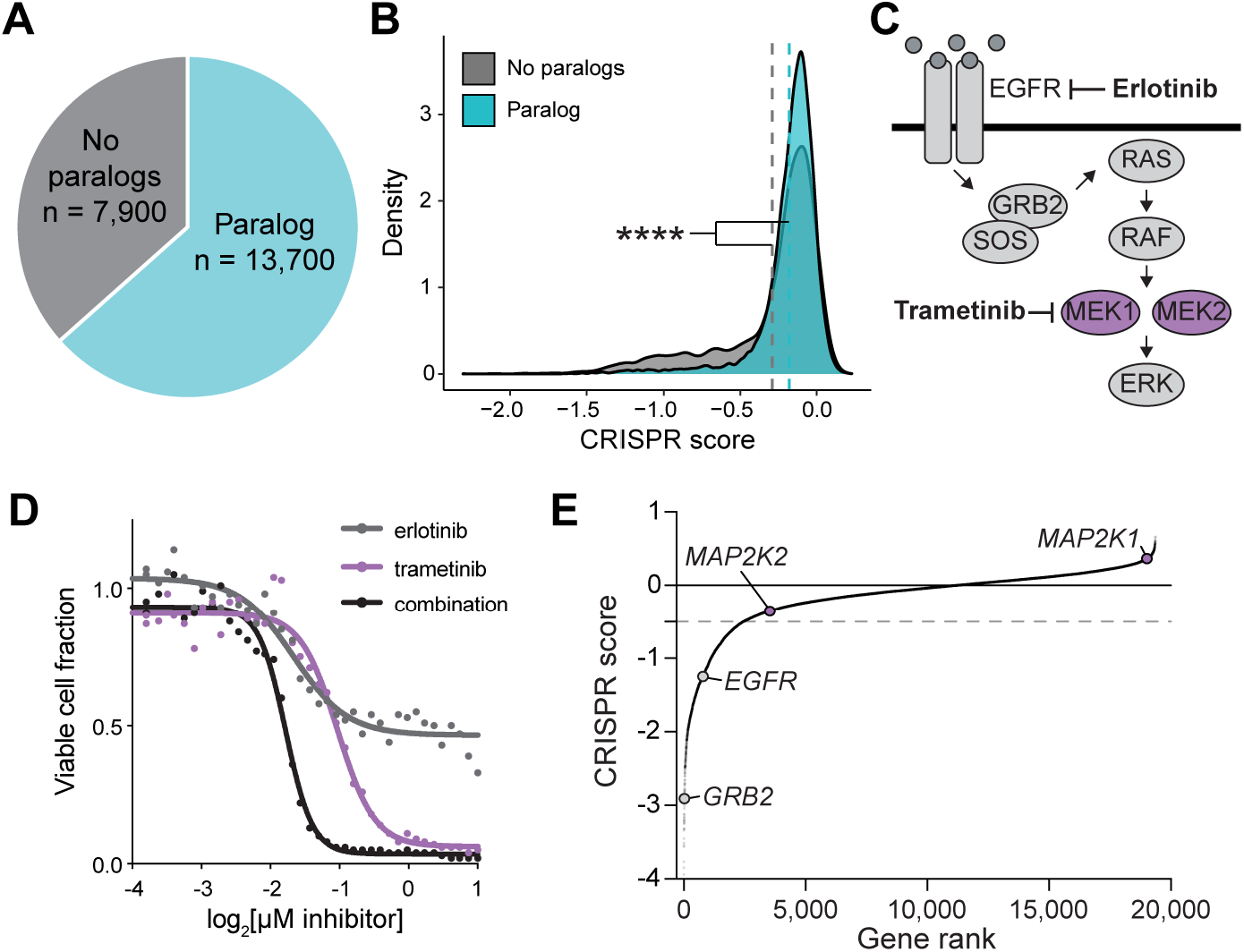
Paralog dependencies are missed in single gene CRISPR knockout screens. **(A)** Pie chart of human genes classified based on whether they are part of a paralog gene family with 50-99% amino acid sequence identity. **(B)** Density plot of CRISPR scores for a single gene CRISPR knockout screen in PC9 lung adenocarcinoma cells. Data is from (Vichas et al., 2020). Dashed lines indicate the mean CRISPR score of genes in each group. *P* < 2.2e-16 by one-tailed Kolmogorov-Smirnov test. **(C)** Schematic of the EGFR receptor signaling pathway. **(D)** Dose response curve of PC9-Cas9-EGFR^T790M/L858R^ lung adenocarcinoma cells treated with erlotinib, trametinib, or a 1:1 combination of both drugs. The fraction of viable cells was determined by CellTiterGlo luminescence after 96 hours of treatment. Data was re-analyzed from a larger drug screen in (Berger et al., 2016). **(E)** Rank plot of CRISPR scores from an erlotinib sensitization screen in PC9-Cas9-EGFR^T790M/L858R^ cells (Vichas et al., 2020). *MAP2K1* (encodes MEK1) or *MAP2K2* (encodes MEK2) single gene knockout does not result in significantly decreased cell growth. Gray dashed lines indicate the threshold for negative selection (−0.5).

To determine whether known therapeutic vulnerabilities are missed in CRISPR knockout screens, we compared our previous drug sensitivity profiling of PC9-EGFR^L858R/T790M^ cells (Berger et al., 2016) to recent genetic vulnerabilities identified in the same system (Vichas et al., 2020). These cells exhibit resistance to the EGFR tyrosine kinase inhibitor, erlotinib, which can be reversed by treatment with trametinib, a kinase inhibitor of MEK1 and MEK2 — protein kinases encoded by the paralogous genes *MAP2K1* and *MAP2K2*, which are part of the Ras/MAPK pathway (**Fig. 1C-D**). The Ras/MAPK pathway is frequently activated in lung cancer by mutation of upstream receptor tyrosine kinases such as EGFR, or activation of KRAS or mutation of *MAP2K1* itself (Arcila et al., 2015; Sanchez-Vega et al., 2018; TCGA, 2014). We noted in singlegene CRISPR knockout data in the same cellular context that while *EGFR* and other Ras pathway members such as *GRB2* were essential as expected, neither *MAP2K1* nor *MAP2K2* was essential when knocked out individually (**Fig. 1E**). We reasoned that paralog redundancy might underlie the apparent disconnect between the small molecule and genetic assays. We therefore sought to develop a multiplexed CRISPR approach to directly probe paralog compensation on a genome scale, enabling the discovery of many more paralogous drug targets that may be missed in current CRISPR-based target discovery efforts.

### The pgPEN library enables single and double knockout of 1,030 human paralog families

To identify synthetic lethal paralogs that could serve as potential lung cancer drug targets, we focused on duplicated genes — paralog families of only two genes. We identified paralog families from Ensembl (Vilella et al., 2009) and then selected families in which a maximum of two genes shared 50-99% amino acid identity (**Supplemental Fig. 1A**). Next, we designed a paired guide RNA (pgRNA) CRISPR library to knock out each paralog alone and in combination with its respective pair. Using pre-validated single guide RNA (sgRNA) sequences from the Brunello CRISPR library (Doench et al., 2016), we designed sixteen four-by-four pairwise double knockout (dKO) pgRNAs for each paralog pair. In addition, we designed single knockout (sKO) pgRNAs containing one targeting sgRNA paired with a non-targeting control sgRNA having no match to the human reference genome. This was done for both paralogs to generate a total of 16 sKO pgRNAs. 500 double non-targeting pgRNAs were included as a control. This “<u>p</u>aired <u>g</u>uide RNAs for <u>P</u>aralog g<u>EN</u>etic interaction mapping (pgPEN)” library (**Supplemental Table 1**) was synthesized and cloned at 1000-fold coverage using previously-developed methods (Gasperini et al., 2017; Thomas et al., 2020). Next-generation sequencing confirmed that >99.99% of pgRNAs were present in the cloned plasmid pool. The final pgPEN library consists of 33,170 pgRNAs targeting 1,030 paralog pairs (2,060 genes) in single knockout and double knockout combinations. Two thirds of paralogs in the pgPEN library are unique to this study while the remainder were also assayed in recent genetic interaction maps (Dede et al., 2020; Gonatopoulos-Pournatzis et al., 2020) (**Supplemental Fig. 1B**). 554 of the gene products of pgPEN-targeted genes are considered “druggable” by recent criteria (Finan et al., 2017) (**Supplemental Fig. 1C**).

To map genetic interactions between paralogs, we applied the pgPEN library to PC9 lung adenocarcinoma cells previously engineered to constitutively express Cas9 (Thomas et al., 2020; Vichas et al., 2020) using standard pooled CRISPR screening methodology in triplicate (**Fig. 2A**). pgRNAs that were positively or negatively selected were identified by Illumina sequencing of pgRNA abundance after ~12 population doublings *in vitro* compared to the starting abundance in the plasmid pool (**Fig. 2A**). Plasmid pgRNA abundance was highly correlated with early time point samples taken immediately following lentiviral transduction and puromycin selection (mean Pearson’s *r* = 0.93; **Supplemental Fig. 1D**). End point samples exhibited expected changes in pgRNA abundance (**Supplemental Fig. 1E**) that were highly correlated across replicates (mean Pearson’s *r* = 0.82; **Supplemental Fig. 1F**). sKO pgRNAs targeting pan-essential genes (Meyers et al., 2017) showed the expected dropout in late time point samples (**Fig. 2B**). These data indicate that the screen was performed without significant bottlenecking and that the pgRNAs performed as expected for known essential genes. Similar to previously established CRISPR screen analysis methods (Meyers et al., 2017), we generated normalized CRISPR scores by scaling pgRNA log2(fold change) values such that the median CRISPR score of double nontargeting constructs was 0 and the median CRISPR score of pan-essential sKO constructs was −1.

**Figure 2.**
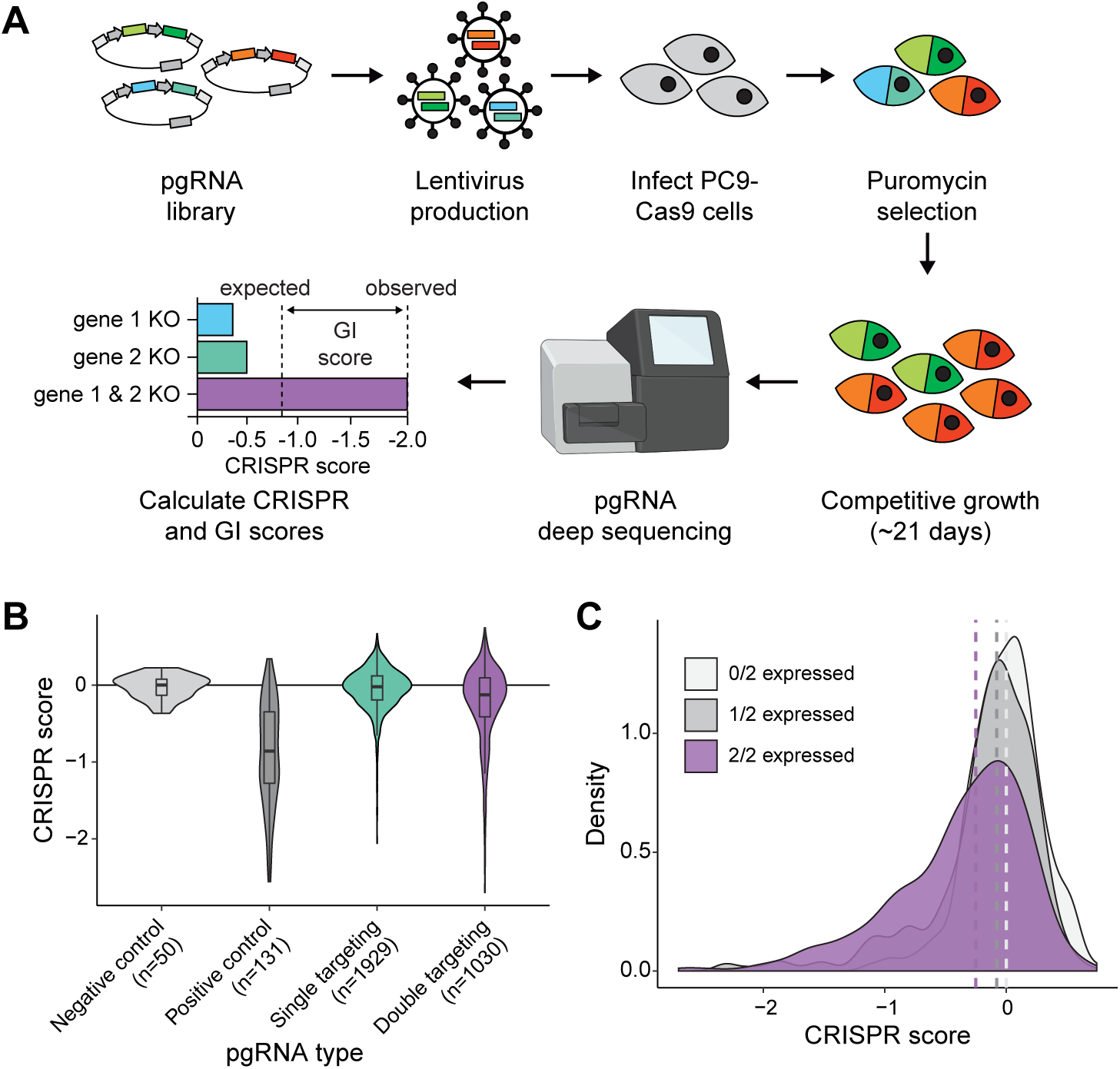
The pgPEN CRISPR library enables genetic interaction mapping of 1,030 human paralog pairs. **(A)** Schematic of pgPEN screening approach for paralog genetic interaction mapping. **(B)** Violin plots of target-level CRISPR scores for negative control (double non-targeting control), positive control (sKO pgRNAs targeting known essential genes), all other sKO, and dKO pgRNAs in the PC9 screen. **(C)** Density plot of target-level CRISPR scores for double targeting pgRNAs grouped by gene expression in PC9 cells indicating for each paralog pair whether zero, one, or both targeted genes are expressed (TPM > 2). Dashed lines indicated the median CRISPR score for each group.

CRISPR-Cas9 gene knockout involves the generation of double strand breaks (DSBs) that can themselves inhibit cell proliferation rate (Aguirre et al., 2016). One concern in targeting multiple loci with Cas9 is that the increased generation of DSBs could, independent of any specific gene effect, result in enhanced negative selection of dKO compared to sKO pgRNAs. To control for this possibility, we further normalized data such that the median CRISPR score of all sKO pgRNAs targeting non-expressed genes would be 0 and the median CRISPR score of all dKO pgRNAs targeting two non-expressed genes would be 0 (Methods and **Supplemental Fig. 1F-G**). After this normalization, dKO constructs still had significantly lower CRISPR scores than sKO constructs *(P* = 1.25e-13 by one-tailed Kolmogorov-Smirnov [K-S] test), indicative of possible genetic interactions in the dKO group. As expected, expressed genes had significantly lower CRISPR scores than unexpressed genes in both single-targeting (*P* < 2.2e-16 by one-tailed K-S test; **Supplemental Fig. 1H**) and double-targeting (*P* < 2.2e-16 by one-tailed K-S test; **Fig. 2C**) constructs. After normalization, only a minimal effect of paralog copy number (**Supplemental Fig. 1I**) on CRISPR score was observed (**Supplemental Fig. 1J-K**). Scaled CRISPR scores for pgRNAs in the PC9 screen can be found in **Supplemental Table 2**.

### Direct identification of paralog genetic interactions in human lung cancer cells

Using the PC9 CRISPR scores, we calculated genetic interaction (GI) scores for each paralog pair under a multiplicative model of genetic interaction following recently developed methods for human GI mapping (DeWeirdt et al., 2020; Han et al., 2017) (Methods). Comparison of the expected and observed CRISPR scores for each paralog pair enabled identification of interacting paralogs (**Fig. 3A**) and calculation of GI scores for each paralog pair (**Fig. 3B-C and Supplemental Table 3**). This approach identified 87 synthetic lethal and 68 buffering genetic interactions among the 1,030 paralog pairs. Synthetic lethal interactions (GI < −0.5 and FDR < 0.1) included *CCNL1/CCNL2, CDK4/CDK6, GSK3A/GSK3B, G3BP1/G3BP2*, and *CNOT7/CNOT8*, as well as *MAP2K1/MAP2K2* (**Fig. 3B-D**), confirming that the discrepancy between genetic and drug data in Figure 1 was indeed due to paralog redundancy. Known synthetic lethal paralogs such as *ARID1A/ARID1B* (Helming et al., 2014) and *MAPK1/MAPK3* (Dede et al., 2020; DeWeirdt et al., 2020) were also identified (**Supplemental Table 3**).

**Figure 3.**
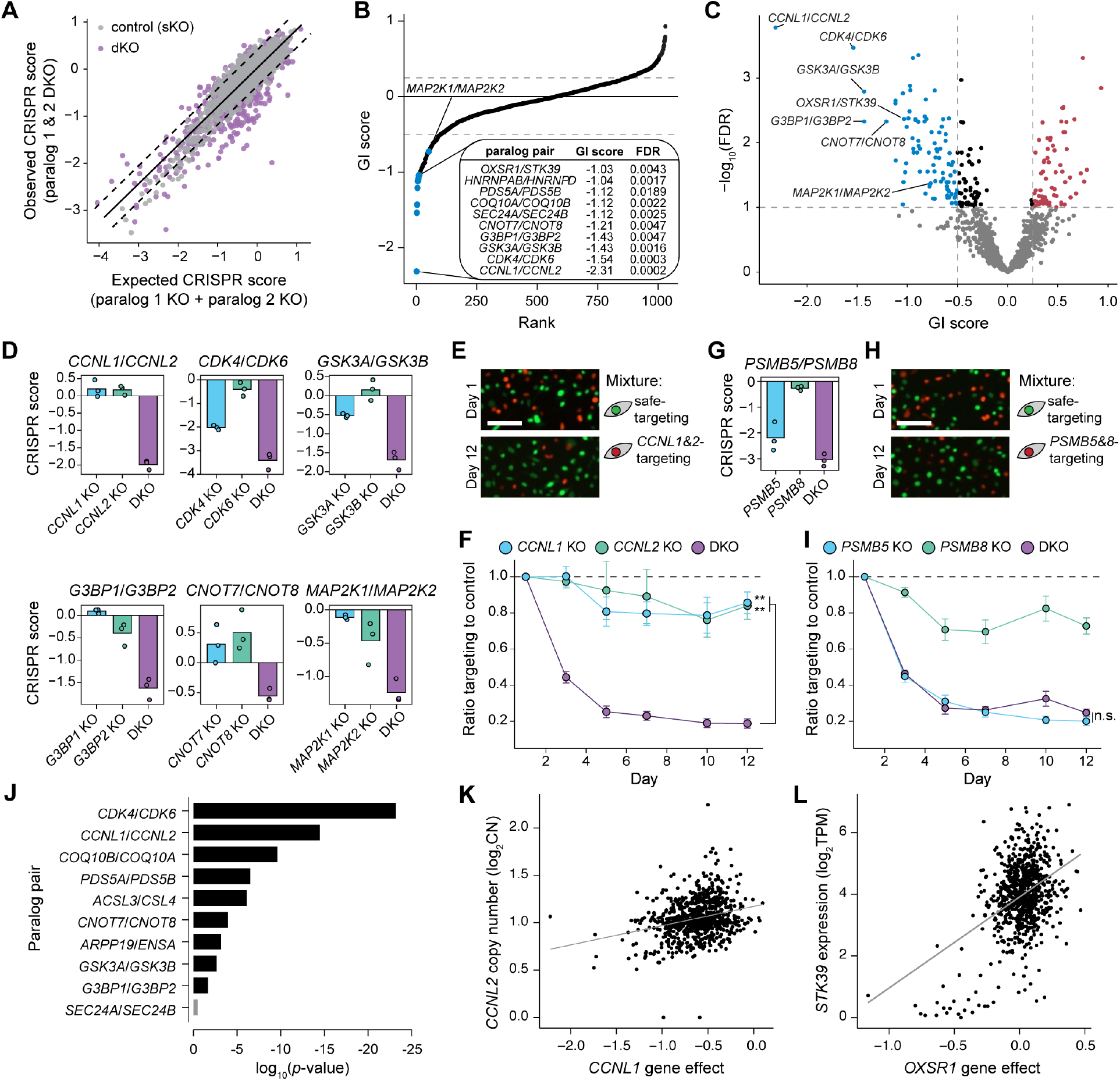
pgPEN uncovers synthetic lethal and buffering genetic interactions. **(A)** Scatter plot of target-level observed versus expected CRISPR scores in the PC9 screen. Dashed lines represents two residuals above or below the linear regression line for the negative control (sKO) pgRNAs. **(B)** Rank plot of target-level genetic interaction scores in PC9 cells. Table insert, top 10 paralogs based on GI score. **(C)** Volcano plot of target-level genetic interaction scores in PC9 cells. FDR indicates the multiple hypothesis-adjusted *P* values from a two-tailed *t*-test (Methods). Blue, synthetic lethal paralog genetic interactions with GI < −0.5 and FDR < 0.1; red, buffering paralog genetic interactions with GI > 0.25 and FDR < 0.1. **(D)** CRISPR scores for representative synthetic lethal paralog pairs. Data shown is the mean CS for each sKO or dKO target across three biological replicates with replicate data shown in overlaid points. **(E)** Fluorescence microscopy images of *CCNL1/CCNL2* competitive fitness assay on Day 0 and Day 12. Scale bar, 100 μM. **(F)** Line graph of competitive fitness assay time course for sKO and dKO of the paralog pair *CCNL1* and *CCNL2.* Data shown is the mean and S.E.M. of six biological replicates. sKO pgRNAs contained one gene-targeting sgRNA and one safe-targeting sgRNA. ** *P* < 8.3e-4 by one-tailed *t*-test. **(G)** CRISPR scores for *PSMB5/PSMB8* in the pooled screen format. Data shown as in (D). **(H)** Fluorescence microscopy images of *PSMB5/PSMB8* competitive fitness assay on Day 0 and Day 12. Scale bar, 100 μM. **(I)** Line graph of competitive fitness assay time course for the paralog pair *PSMB5* and *PSMB8*, performed as in (F). n.s., not significant by one-tailed *t*-test. **(J)** Bar plot indicating the best linear regression *P* value obtained by comparing the gene effect of a single paralog knockout to the expression or copy number of its pair across cell lines in DepMap single-gene CRISPR knockout screen data (www.DepMap.org). Black, *P* < 0.05; gray, *P* ≥ 0.05. **(K)** Scatter plot comparing the effect of CRISPR-mediated knockout of *CCNL1* to the copy number of its paralog *CCNL2.* Each dot represents a cell line. Data retrieved from DepMap.org. **(L)** Scatter plot comparing the effect of CRISPR-mediated knockout of *OXSR1* to the expression of its paralog *STK39.* Each dot represents a cell line. Data retrieved from Depmap.org.

To experimentally validate these findings, we developed a competitive growth assay in red (mCherry) and green (GFP) labeled PC9-Cas9 cells (**Supplemental Fig. 2**). In designing this validation experiment, we used safe-targeting gRNAs (Morgens et al., 2017) in place of nontargeting gRNAs to account for the growth effects observed by generating one versus two doublestrand breaks. We transduced PC9-Cas9-GFP-NLS cells with a double safe-targeting pgRNA, while PC9-Cas9-mCherry-NLS cells were transduced with paralog-targeting pgRNAs designed to knock out the expression of each paralog individually or both paralogs together (**Supplemental Table 4**). Double safe-targeting cells were pooled with paralog-targeting cells at a 1:1 ratio of GFP:mCherry cells. Using this approach, we determined the effect of one synthetic lethal paralog pair, *CCNL1/CCNL2*, and one non-synthetic lethal paralog pair, *PSMB5/PSMB8.* The results of these competition assays mirrored the gene knockout effects observed in the pooled screen format (**Fig. 3E-I**). For *CCNL1/CCNL2*, individual gene knockouts showed little effect on cell growth, whereas combined knockout of *CCNL1* and *CCNL2* resulted in severe growth effects in both the screen (**Fig. 3E**) and the competition assay (*P* = 1.32e-05 for *CCNL1* sKO vs. dKO, *P* = 4.12e-4 for *CCNL2* sKO vs. dKO by one-tailed *t*-test; **Fig. 3F**). In contrast, *PSMB5/PSMB8* combined knockout was not significantly different from *PSMB5* single knockout in the PC9 CRISPR screen (**Fig. 3G**), consistent with validation experiments in the competitive growth assay (*P* = 0.91 by one-tailed *t*-test; **Fig. 3H-I**).

As a complementary approach to validate our screen, we used single gene knockout data from DepMap.org (Tsherniak et al., 2017) to determine whether other identified synthetic lethal paralog pairs showed synthetic lethality in the context of spontaneous loss of one member in cancer cell lines. Of the top ten synthetic lethal pairs identified in the screen, 9/10 showed a significant association between a single gene knockout “gene effect” and either expression or copy number of the other paralog (**Fig. 3J** and **Supplemental Table 5**). These included *CCNL1/CCNL2*, genes involved in pre-mRNA splicing of many transcripts including apoptotic genes, (**Fig. 3K**) and *OXSR1/STK39*, which encode kinases involved in the oxidative stress response (**Fig. 3L**). Others such as *SEC24A/SEC24B* did not show any significant associations in the DepMap analysis, prompting us to investigate whether there are tissue-specific essential paralog families that may be missed by broad analyses of multiple tissues such as DepMap data in aggregate.

### A second pgPEN screen identifies conserved versus tissue-specific paralog synthetic lethal interactions

Next, we applied the pgPEN approach to a different tissue context, HeLa cervical carcinoma cells, using similar methodology with the exception of using a doxycycline-inducible Cas9 system (Cao et al., 2016). Quality control analyses of HeLa screening data again indicated successful generation of expected gene knockout phenotypes (**Supplemental Fig. 3A-J** and **Supplemental Table 6**). Calculation of GI scores identified 70 significant synthetic lethal interactions and 44 significant buffering interactions (**Fig. 4A-B**, **Supplemental Fig. 3K**, and **Supplemental Table 4**). Many of the top synthetic lethal pairs were shared between HeLa and PC9 cells, including *CCNL1/CCNL2, GSK3A/GSK3B*, and *MAP2K1/MAP2K2* (**Fig. 4C-D**). Other paralog families were synthetic lethal only in one of the cell contexts. These included *SOS1/SOS2*, which were highly essential and synthetic lethal only in HeLa cells (**Fig. 4C-D**) and *CDK4/CDK6*, which were only required in PC9 cells (**Fig. 4C-D**). In total, 122 paralog pairs were identified as synthetic lethal in at least one context. Surprisingly, we noted that cell-type specific synthetic lethal interactions were often not explained by expression differences (**Fig. 4D**), demonstrating that paralog dependencies, like other cancer dependencies, are modified by cellular context or other biological factors besides gene expression.

**Figure 4.**
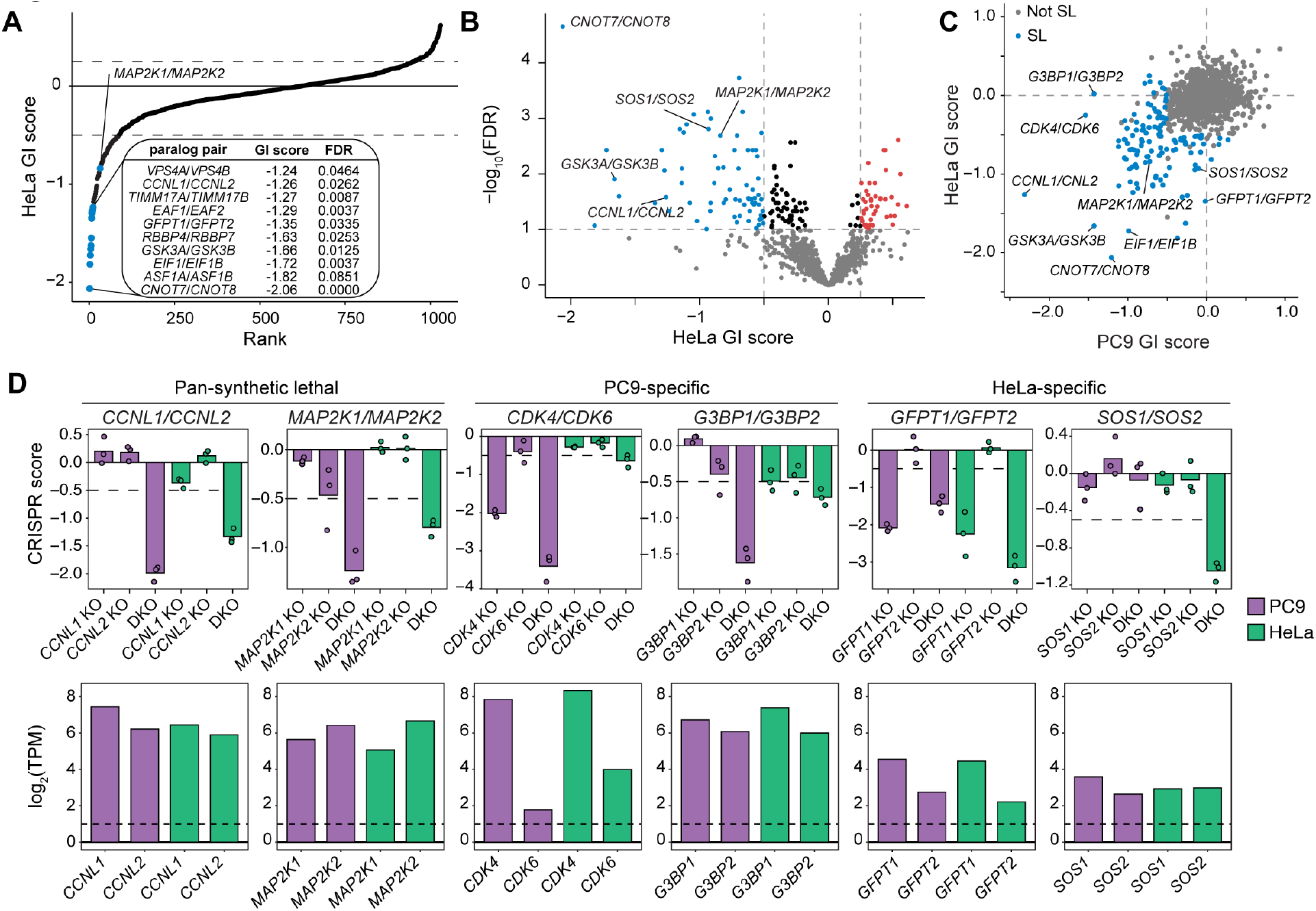
Identification of context-specific and pan-essential synthetic lethal paralog pairs. **(A)** Rank plot of target-level genetic interaction scores in HeLa cells. **(B)** Volcano plot of target-level genetic interaction scores in HeLa cells. FDR indicates the multiple hypothesis-adjusted *P* values from a two-tailed *t*-test (Methods). Blue, synthetic lethal paralog genetic interactions with GI < −0.5 and FDR < 0.1; red, buffering paralog genetic interactions with GI > 0.25 and FDR < 0.1. **(C)** Scatter plot of target-level genetic interaction scores for paralog pairs in PC9 versus HeLa cells. Blue, synthetic lethal paralog pairs with GI < −0.5 and FDR < 0.1 in either PC9 or HeLa cells; gray, all paralog pairs with GI ≥ −0.5 or FDR ≥ 0.1. **(D)** CRISPR scores for representative synthetic lethal paralog pairs identified in the PC9 and HeLa cell screens. Top row: Data shown is the mean CS for each sKO or dKO target across three biological replicates with replicate data shown in overlaid points. Pan-synthetic lethal paralogs have FDR < 0.1 in both cell lines, PC9-specific paralogs have FDR < 0.1 in PC9 only, and HeLa-specific paralogs have FDR < 0.1 in HeLa only. Bottom row: Paralog gene expression in PC9 and HeLa cells from RNA-seq analysis.

Some synthetic lethal paralog pairs, including *SEC24A/SEC24B, COQ10A/COQ10B, CNOT7/CNOT8, TIA/TIAL*, and *VPS4/VPS4B* have been highlighted in previous studies (Dede et al., 2020; Gonatopoulos-Pournatzis et al., 2020; Lord et al., 2020; Neggers et al., 2020; Szymańska et al., 2020). However, to our knowledge many of the synthetic lethal paralogs identified in the pgPEN screens were not previously known to be functionally redundant in human cells. These include *CCNL1/CCNL2* and *OXSR1/STK39* along with eukaryotic translation initiation factors *EIF1/EIF1B*, DNA and RNA helicase and cGAS/STING pathway members *G3BP1/G3BP2*, hexosamine biosynthesis pathway members *GFPT1/GFPT2*, and *PDS5A/PDS5B*, which regulate sister chromatid cohesion during mitosis. Individual members of many of these novel synthetic lethal paralog families have been previously implicated in cancer; for instance, high *GFPT2* expression has been linked to tumor metabolic reprogramming in lung adenocarcinoma (Zhang et al., 2018) and *PDS5B* is a negative regulator of cell proliferation and has been highlighted as a possible tumor suppressor gene in prostate cancer (Maffini et al., 2008).

### Identification of tumor suppressor paralog pairs

In addition to synthetic lethal interactions, pgPEN screening can identify positive genetic interactions. We noticed that these positive interactions include both buffering interactions, where loss of one paralog prevents the deleterious phenotype of loss of the other — we identified 108 of such interactions in at least one cell context — as well as cases where combined loss of both genes synergistically promote cell growth. The latter are likely to be paralog families with tumor suppressive functions that require complete loss of the family to reveal the cellular phenotype. To identify these tumor suppressive paralogs, we restricted our analysis to significant buffering interactions (GI > 0.25, FDR < 0.1) between expressed paralogs in which the double knockout was positively enriched in each CRISPR screen (CS > 0.25). Under these relatively stringent criteria, four tumor suppressor buffering interactions were identified in PC9, and six in HeLa cells (**Fig. 5A**). None of the ten interactions were shared across cell lines, potentially reflecting the differing biology of HeLa and PC9 cells and the difficulty in achieving positive selection in basal culture conditions of rapidly proliferating cancer cell lines.

**Figure 5.**
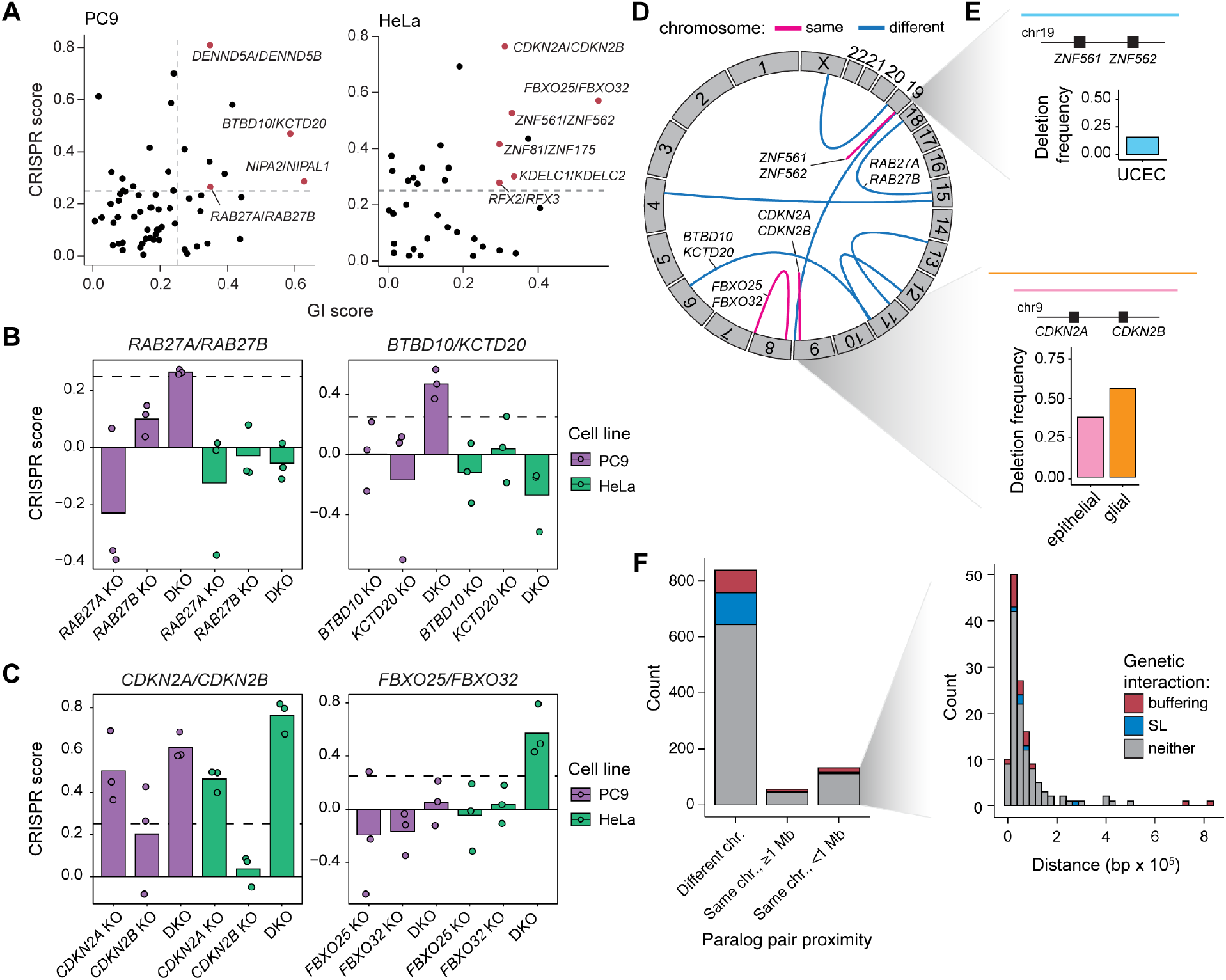
Paralog buffering interactions include tumor suppressor paralogs. **(A)** Identification of tumor suppressive paralog interactions (GI score > 0.25; FDR < 0.1; CRISPR score > 0.25). **(B)** CRISPR scores of PC9-specific tumor suppressive paralog pairs. Data shown is the mean of three biological replicates with replicate data shown in overlaid points. **(C)** CRISPR scores of HeLa-specific tumor suppressive paralog pairs. Data shown as in (B). **(D)** Circos plot showing the genomic locations of tumor suppressive paralog pairs. Blue arcs indicate paralog pairs located on different chromosomes, while pink arcs represent paralog pairs located on the same chromosome. **(E)** Top, diagram of a recurrent deletion seen in uterine corpus endometrial carcinoma (UCEC) data from TCGA that spans the genomic locus containing *ZNF561* and *ZNF562*, and a bar plot indicating the deletion frequency. Bottom, diagram of recurrent deletions in epithelial and glial cancers that span the genomic locus containing *CDKN2A* and *CDKN2B*, and a bar plot showing the deletion frequency in each cancer subtype. **(F)** Genomic distance between paralogs for the 1,030 paralog pairs included in the pgPEN library in three proximity categories: on different chromosomes, on the same chromosome but ≥1 megabase apart, and on the same chromosome within 1 megabase. Inset: Histogram of paralog distance for pairs that are within 1 megabase of one another.

Tumor suppressor pairs identified in PC9 cells include *RAB27A/RAB27B*, encoding Rab-family GTPases involved in vesicle trafficking (Li et al., 2018), and the BTB/POZ-domain genes *BTBD10/KCTD20* (**Fig. 5B**). In HeLa cells, one of the top pairs identified was *CDKN2A/CDK2NB* (**Fig. 5C**), frequently deleted tumor suppressors that encode the CDK4/6 inhibitors INK4A/ARF and INK4B (Kim and Sharpless, 2006). A novel top tumor suppressor paralog pair in HeLa cells was *FBXO25/FBXO32* (**Fig. 5C)**, which encode SCF-type E3 ligase proteins. Although little is known about the function of these proteins and their substrates, the genetic interaction between *FBXO25* and *FBXO32* suggests that these two proteins may share similar functions or substrates. *FBXO25* and *FBXO32* have been individually proposed to have tumor suppressive function in previous studies (Xue et al., 2012; Zhou et al., 2017).

While this direct identification of tumor suppressor paralog pairs has merit for understanding basic genome function, spontaneous loss of two unlinked genes in cancer should be rare, and therefore it is unlikely that double knockout of paralogs contributes to tumorigenesis for most paralog families. Interestingly however, we noted that two tumor suppressor pairs contained genes co-located in the same chromosomal locus (**Fig. 5D**). In addition to *CDKN2A/CDKN2B*, whose combined loss is well known to promote tumorigenesis, *ZNF561* and *ZNF562* are also co-located and reside on chromosome 19p13.2, a frequently deleted region in uterine corpus endometrial cancer (Berger et al., 2018; Cherniack et al., 2017) (**Fig. 5E**). Beyond the tumor suppressor paralogs, 13% of all the paralog pairs in the pgPEN library are located within 1 MB of each other in the human genome (**Fig. 5F**), which raises the possibility that the cell fitness consequences of double knockout of human paralogs could contribute to the selective forces that drive aneuploidy patterns in human cancer (Ben-David and Amon, 2020; Taylor et al., 2018).

## Discussion

This work provides, to our knowledge, the largest direct experimental assessment of paralog genetic interactions in the human genome to date. The pgPEN library we developed uses two Cas9-type sgRNAs driven from independent promoters to enable knockout of two paralogs simultaneously and targets 2,060 duplicate human paralogs. Complementing two other recent studies of human paralog genetic interactions (Dede et al., 2020; Gonatopoulos-Pournatzis et al., 2020), our library adds 1,331 unique paralogs and brings the total set of human paralogs assayed to date to 3,237. An additional difference between the present study and these other reports is our use of Cas9 versus the newer Cas12a-derived enzymes. Cas12a systems have the benefit of using an array of sgRNAs on a single transcript that is processed by Cas12a, enabling programmable delivery of multiple sgRNAs to the same cell (DeWeirdt et al., 2020). Continued application of Cas12a for CRISPR screening will enable the experimental identification of higher order combinatorial genetic interactions in human cells. However, for pairwise interactions of paralogs, the pgPEN library may provide an ease of application to investigators with Cas9-expressing cell systems already developed.

Remarkably, we find that 12% of duplicate paralogs exhibit synthetic lethality, demonstrating that paralogs are a rich source of genetic interactions. These findings underscore the importance of simultaneously targeting redundant genes and demonstrate that a large fraction of cancer dependencies are missed by current single-gene knockout approaches. Like others (De Kegel and Ryan, 2019; Viswanathan et al., 2018), we propose that synthetic lethal interactions among paralogs could be harnessed for cancer therapy, since the aneuploid genomes typical of cancer cells commonly harbor deletions and inactivating mutations in one or more paralogs. Targeting lineage-specific essential paralogs or paralog families with partial loss in cancer could provide an orthogonal approach for cancer therapy to be applied in combination with existing therapies to provide durable cancer control and improved patient outcomes. In addition, even paralogs that are not lost in cancer may represent tractable cancer targets; the same homology and redundancy that complicates genetic identification of paralogs as cancer dependencies could enable simultaneous targeting of each protein with ease. This strategy is exemplified by the current use of small molecules targeting several of the top synthetic lethal paralogs we identified, such as *CDK4/CDK6* and *GSK3A/GSK3B.*

Last, we provide the first systematic identification of tumor suppressive paralog pairs. We identified ten paralog pairs whose combined loss significantly promotes cancer cell line growth. Although combined loss of some of these pairs is likely to be rare, 2 of 10 we identified are located in the same chromosomal locus. Many of these loci are frequently deleted in cancer. These data therefore shed light on the basis for the positive selection of these genome deletions and suggest that combined paralog loss may shape the landscape of positive and negative selection in human cancer.

### Methods

#### Human paralog analysis and selection

For analysis of human paralog versus singleton essentiality (**Fig. 1A-B**), a list of human proteincoding genes was obtained from Ensembl. Mitochondrial genes and splice variants were removed from the analysis. The remaining genes were divided into two groups: (1) paralogous genes with >10% amino acid sequence identity and (2) singleton genes.

For the pgPEN library, the list of human paralogs was further filtered to include only those with >50% reciprocal amino acid sequence identity with only one other gene. Genes encoding components of olfactory signaling and T cell receptors were also excluded. As shown in **Supplemental Figure 1A**, a total of 2,060 paralogous genes (1,030 pairs) was included in the pgPEN library.

#### PC9 single-gene CRISPR knockout screen and drug sensitivity profiling data

PC9-Cas9 and PC9-Cas9-EGFR^T790M/L858R^ single-gene CRISPR knockout screen data (Vichas et al., 2020) and erlotinib/trametinib drug sensitivity data (Berger et al., 2016) were re-analyzed from previously published data. The relative essentiality of singletons versus paralogs in the PC9-Cas9 CRISPR knockout screen using the Brunello library (Doench et al., 2016) was assessed via a two-tailed Kolmogorov-Smirnov test (**Fig. 1B**). Drug sensitivity data was generated as part of a large-scale screen in (Berger et al., 2016) performed in PC9 cells ectopically expressing the erlotinib-resistance variant, EGFR^T790M/L858R^. For combination dosing, erlotinib and trametinib were delivered to cells in a 1:1 molar ratio.

#### Paralog pgRNA plasmid library design and cloning

The pgPEN library was designed using pre-validated sgRNAs selected from the Brunello library (Doench et al., 2016). sgRNAs containing BsmBI restriction target sequences and U6 termination signals were excluded from the library. Given that previous data demonstrated no position effects using the pgRNA approach (Gasperini et al., 2017), the sgRNA targeting a given gene was located at the same site in every pgRNA.

pgRNA oligonucleotides were synthesized by Twist Biosciences and cloned per published protocols (Gasperini et al., 2017; Thomas et al., 2020). Briefly, the pgRNA oligonucleotides were amplified (primers RKB1169 and RKB1170, **Supplemental Table 7**) using NEBNext High Fidelity 2X Ready Mix (New England Biolabs) and purified via a 1.8X Ampure XP SPRI bead (Beckman Coulter) clean-up. Amplified oligonucleotides were then cloned into BsmBI (FastDigest Esp3I, Thermo Fisher Scientific)-digested lentiGuide-Puro (Addgene no. 52963) plasmid backbone via the NEBuilder HiFi (New England Biolabs) assembly system. Cloned plasmids were purified using a 0.8X Ampure XP SPRI bead clean-up and transformed into Endura ElectroCompetent *E. coli* cells (Lucigen) via electroporation to generate the pLGP-2xSpacer vector. The pLGP-2xSpacer vector was isolated using the NucleoBond Xtra Maxiprep kit (Macherey-Nagel) and linearized by BsmBI digest. A GBlock (synthesized by Integrated DNA Technologies) containing a second gRNA backbone and H1 promoter sequence was digested with BsmBI, purified via a 1.8X Ampure bead clean-up, and ligated into the pLGP-2xSpacer backbone using NEB Quick Ligase (New England Biolabs). The reaction product was purified using an 0.8X Ampure bead cleanup and transformed into Endura Electrocompetent *E. coli* via electroporation to propagate the final pLGP-pgRNA vectors. The pLGP-pgRNA plasmids were again isolated using the NucleoBond Xtra Maxiprep kit, and the cloned library was amplified and sequenced as described below to confirm high coverage. At each cloning steps, individual *E. coli* colonies were sequence verified via colony PCR and Sanger sequencing with primer RKB1148. Over 1000X coverage of each pgRNA was maintained throughout plasmid library cloning, amplification, and sequencing; coverage depth was selected based on our previous screen experience as well as published recommendations (Doench, 2018; Joung et al., 2017).

#### Lentivirus production and titration

With our cloned library, we produced lentivirus via a large-format transfection in HEK293T cells using a protocol adapted from Joung *et al.* (Joung et al., 2017). Briefly, we used TransIT-LT1 (Mirus Bio) as a transfection reagent, with packaging plasmid psPAX2 (Addgene no. 12260) and envelope plasmid pVSV-G (Addgene no. 8454) and Opti-MEM (Thermo Fisher Scientific). Plasmids were added at a 4:2:1 ratio of transfer to packaging to envelope plasmid. 18 hours post-transfection, media was changed to high-serum DMEM (30% FBS). Lentivirus was harvested 48 hours post-transfection. Over 500X coverage of each pgRNA was maintained throughout; coverage depth was selected based on our previous screen experience as well as published recommendations (Doench, 2018; Joung et al., 2017).

#### Cell lines

PC9-Cas9 cells were a gift from Dr. Matthew Meyerson (Broad Institute) and cultured in RPMI-1640 (Gibco) supplemented with 10% Fetal Bovine Serum (FBS, Sigma). PC9-Cas9-GFP-NLS and PC9-Cas9-mCherry-NLS cells were generated by transducing PC9-Cas9 cells with lentivirus containing GFP-NLS or mCherry-NLS-encoding vectors (parental backbone, Addgene no. 12252). mCherry or GFP positive cells were selected using flow cytometry. HeLa/iCas9 cells were previously generated (Cao et al., 2016) and cultured in Dulbecco’s Modified Eagle’s Medium (DMEM, Genesee Scientific) supplemented with 10% FBS. All cells were maintained at 37°C in 5% CO2 and confirmed mycoplasma-free.

#### Genome-wide CRISPR paralog knockout screen

PC9-Cas9 and iCas9/HeLa cells were transduced with the pgPEN library at low multiplicity of infection (~0.3) to ensure the integration of a single pgRNA construct into >95% of cells (Doench, 2018). Transduced cells were then selected using puromycin (1.0 μg/mL, Sigma) for 48-72 hours until all uninfected control cells were dead. For the PC9-Cas9 screen, cells were split into three biological replicates after infection but before puromycin selection, and genomic DNA (gDNA) was harvested from each replicate after puromycin selection for an early time point sample. For the iCas9/HeLa screen, cells were kept in the pooled format until puromycin selection was complete, resulting in a single early time point sample. iCas9/HeLa cells were then induced using doxycycline (1.0 μg/mL, Sigma) and split into three biological replicates. For both screens, cells were then passaged for approximately 12 population doublings while maintaining over 500X coverage of each pgRNA at every step. An endpoint gDNA sample was harvested from each biological replicate and stored at −80°C. Genomic DNA was extracted using the QIAamp DNA Blood Maxi Kit (Qiagen).

#### Paralog pgRNA amplification, library preparation, and sequencing

Plasmid and gDNA samples were amplified and sequenced at >500X coverage per pgRNA as per our previously established methods (Thomas et al., 2020). First, 2.5 μg of gDNA was used as input for each reaction, with a total of 48 reactions (120 μg total input gDNA) to ensure >500X coverage per sample. Input DNA was amplified using NEBNext High Fidelity 2X Ready Mix with primers RKB2713/RKB2714 followed by 1.8X Ampure bead clean-up. Second, the amplicon from PCR no. 1 was used as input for PCR no. 2, with 10 ng input DNA in one reaction per sample. The input DNA was amplified using primers RKB2715/RKB2716 followed by 1X Ampure bead clean-up. Third, 10 ng of the amplicon from PCR no. 2 was used as input for PCR no. 3 and was amplified using a common forward primer (RKB2717) and a sample-specific barcoded reverse primer (see **Supplemental Table 7**) to allow for multiplexed sequencing. Product from PCR no. 3 was purified using a 1X Ampure bead clean-up, quantified by a Qubit assay (Thermo Fisher Scientific), and pooled at equimolar amounts prior to Illumina sequencing.

#### Paralog pgRNA CRISPR screen analysis

Sequencing reads for each pgRNA were mapped separately to the pgPEN library annotation using Bowtie (Langmead et al., 2009). Based on the reference set, correctly-paired pgRNAs were retained while incorrectly-paired gRNAs were discarded. pgRNAs with <2 reads per million (RPM) in the plasmid pool or with a read count of zero at any time point were also removed. The log2-scaled fold change (LFC) of each pgRNA was then computed using MAGeCK (Li et al., 2014) to compare initial abundance in the plasmid pool to abundance at early and late time points.

LFC values were scaled so that the median of negative control (double non-targeting) pgRNAs was set to zero, while the median of positive control (single-targeting pgRNAs targeting Project Achilles pan-essential genes (Meyers et al., 2017) pgRNAs was set to −1.0 (**Fig. 2B** and **Supplemental Fig. 3D**). We also used RNA-seq data from each cell line (Thomas et al., 2020) to control for growth defects caused by the double-strand break generation and repair process. To do this, we adjusted pgRNA LFCs so that the median LFC of single- and double-targeting gRNAs targeting unexpressed genes (TPM <2) was set to zero (**Fig. 2C** and **Supplemental Figs. 1G-H** and **3E-G**). Finally, we analyzed copy number effects using data from DepMap.org. We grouped pgRNAs by the combined copy number of targeted genes for each construct and analyzed the CRISPR scores of each copy number group (**Supplemental Figs. 1I-K** and **2H-J**). Given that the copy number of the vast majority of paralogs included in our library was close to 2, we did not adjust for copy number effects. The scaled and normalized LFC for each pgRNA was termed a CRISPR score (CS). Target-level CRISPR scores were calculated by taking the mean across pgRNAs with the same sKO or dKO paralog target. Final CRISPR scores were computed by taking the mean across the three biological replicates for each screen.

#### Genetic interaction score calculations

To compute a genetic interaction score for each paralog pair, we combined two previously published methods for genetic interaction mapping in human cells (DeWeirdt et al., 2020; Han et al., 2017). We first calculated an expected and observed CS for each pgRNA. For dKO pgRNAs (pgRNA-Paralog1_Paralog2), we calculated the expected CS by first taking the mean CRISPR scores of each sKO pgRNA with the same targeting sgRNA sequence paired with a NTC sgRNA sequence (i.e., mean(pgRNA-Paralog1_NTC1, pgRNA-Paralog2_NTC2) and mean(pgRNA-NTC1_Paralog2, pgRNA-NTC2_Paralog2)). We summed these two sKO mean CS values to calculate an expected CS for each paralog pair, and compared this expected CS to the observed dKO CS (pgRNA-Paralog1_Paralog2). For sKO pgRNAs, we calculated an expected CS by computing the sum of (1) the CS for the other sKO pgRNA containing the same targeting sgRNA sequence paired with a different NTC sgRNA sequence (pgRNA-Paralog1_NTC2) and (2) the mean CS of double NTC pgRNAs (pgRNA-NTC1_NTC2) containing the same NTC sgRNA sequence (i.e., mean(pgRNA-NTC1_NTC2, pgRNA-NTC1_NTC3)). This sKO expected CS was then compared to the observed sKO CS (pgRNA-Paralog1_NTC1 or pgRNA-NTC1_Paralog2). Target-level sKO and dKO expected and observed CRISPR scores were calculated by taking the mean across pgRNAs.

We then obtained the distribution of CRISPR scores for control pgRNAs by calculating the linear regression of control (sKO) expected versus observed CS values (**Fig 3A** for PC9 and **Supplemental Fig 3L** for HeLa). GI scores were calculated by calculating the residual of each observed CS value for each paralog pair from the control regression line. A *t*-test was used to compute the statistical significance of the difference in dKO GI scores versus the distribution of control (sKO) GI scores. A Benjamini-Hochberg false discovery rate (FDR) correction (Benjamini and Hochberg, 1995) was then applied, and FDR < 0.1 was considered significant.

Synthetic lethal paralogs were defined as those with a GI score < −0.5 and FDR < 0.1, while buffering paralogs were defined as those with GI score > 0.25 and FDR < 0.1. Tumor suppressor paralogs were defined as buffering paralogs with an additional filter for dKO CS > 0.25 in either PC9 or HeLa cells. Cancer deletion data for paralog tumor suppressor analysis shown in **Figure 5E** were obtained from The Cancer Genome Atlas Copy Number Portal (Beroukhim et al., 2010).

#### Competition assay

For the bichromatic competition assay, PC9-Cas9-GFP-NLS cells were infected with a control pgRNA and PC9-Cas9-mCherry-NLS cells with either a paralog sKO pgRNA or a paralog dKO pgRNA. After 48-72 hours of selection with puromycin (1 μg/mL), cells were pooled at an equal ratio and seeded in tissue culture-treated plates (Corning). All pgRNA sequences used for the competition assay are available in **Supplemental Table 4**. For the *MAGOH/MAGOHB* paralog pair, NTC gRNAs were used as controls and each competition (double NTC *versus MAGOH* sKO, double NTC *versus MAGOHB* sKO, double NTC *versus MAGOH/MAGOHB* dKO) was carried out in triplicate. For the *CCNL1/CCNL2* and *PSMB5/PSMB8* competition assays, safe-targeting gRNAs that target intergenic regions (Morgens et al., 2017) were used as controls to account for the different number of double-strand breaks generated by sKO versus dKO pgRNAs. Each *CCNL1/CCNL2* and *PSMB5/PSMB8* competition was carried out in six biological replicates. After pooling, cells were imaged and raw counts of mCherry- and GFP-expressing cells were computed every 2-3 days using a Cytation 5 imager (BioTek Instruments). The ratio of mCherry to GFP was computed and normalized to the Day 0 count. At the assay end point, the targeting-to-control ratios were compared for sKO versus dKO conditions via a one-tailed Student’s *t*-test.

#### Statistics and reproducibility

Statistical tests are indicated in the figure legends. Results were analyzed for statistical significance with R v3.6.3 in an Rstudio v1.2.5 environment or in GraphPad Prism 8.

## Data Availability Statement

RNA-seq data for PC9 and HeLa cells are available from GSE120703. The authors declare that all other data supporting findings of this study are available within the paper and its Supplemental information files.

## Acknowledgments

We thank Dr. Athea Vichas (Fred Hutchinson Cancer Research Center) for advice on CRISPR screens in PC9 cells. This work was funded with support from the Lung Cancer Research Foundation. P.C.R.P. was supported by NSF DGE-1762114 and NIH T32-HG000035. J.D.T. is a Washington Research Foundation Postdoctoral Fellow. A.H.B. was supported in part by the NIH/NCI (R00 CA197762); the Devereaux Outstanding Investigator Award from the Prevent Cancer Foundation; the Stephen H. Petersdorf Lung Cancer Research Award; Seattle Translational Tumor Research program; and the Innovators Network Endowed Chair. RKB was supported in part by the NIH/NIDDK (R01 DK103854); NIH/NHLBI (R01 HL128239 and R01 HL151651); NIH/NCI (R01 CA251138); Edward P. Evans Foundation; Blood Cancer Discoveries Grant program through the Leukemia & Lymphoma Society, Mark Foundation for Cancer Research, and Paul G. Allen Frontiers Group (8023-20); Dept. of Defense Breast Cancer Research Program (W81XWH-20-1-0596); and the McIlwain Family Endowed Chair in Data Science. RKB is a Scholar of The Leukemia & Lymphoma Society (1344-18). Sequencing was performed by the Fred Hutch Genomics Shared Resource (supported by NIH/NCI Cancer Center Support Grant P30 CA015704). Computational studies were supported in part by FHCRC’s Scientific Computing Infrastructure (ORIP S10 OD028685).

## Contributions

A.H.B conceived of the project. A.H.B. and R.K.B. supervised the project. P.C.R.P, J.D.T., and A.H.B. designed experiments. P.C.R.P., J.D.T., S.K., and A.G. conducted experiments. P.C.R.P and J.D.T. analyzed the data. P.C.R.P., J.D.T., and A.H.B. wrote the manuscript.

---

**Supplemental Figure 1.**
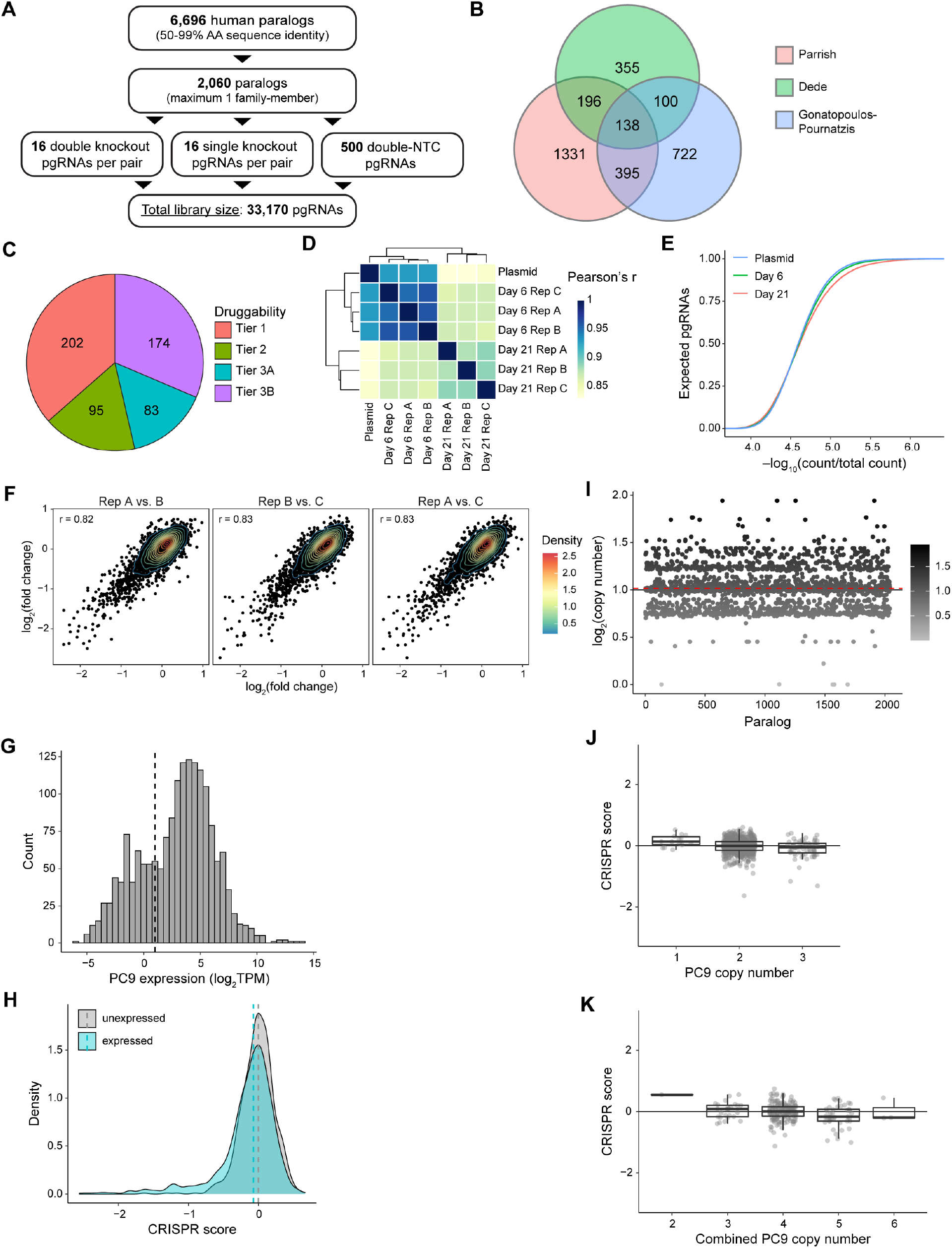
**(A)** Schematic of filtering strategy used to select paralogous genes for inclusion in the pgPEN library. **(B)** Venn diagram of paralogs included in this study, Dede et al. (Dede et al., 2020), and Gonatopoulos-Poumatzis et al. (Gonatopoulos-Pournatzis et al., 2020). **(C)** Pie chart depicting the number of genes considered “druggable” based on classification from Finan et al. (Finan et al., 2017). **(D)** Heat map illustrating Pearson correlations between sequencing reads in counts per million (CPM) supporting each pgRNA for all samples from the PC9 screen. Dendrogram, unsupervised clustering of CPM by the complete-linkage method. **(E)** Plot showing the cumulative distribution of pgRNA counts at each time point (mean across three biological replicates) in the PC9 screen. Each point represents the proportion of total pgRNAs with the corresponding count in a given sample. **(F)** Scatter plot of target-level log2(fold changes) between the plasmid and late time points for the indicated replicate comparisons in the PC9 screen. Contour lines indicate the density of points in 2D space. **(G)** Histogram of PC9 RNA-seq reads in log2(TPM) for all paralogs in the pgPEN library. Dashed line indicates the cutoff used for expression: log2(TPM) > 1. **(H)** Density histogram of CRISPR scores for sKO paralog target genes, grouped by target gene expression in PC9. **(I)** Scatter plot of PC9 log2(copy number) for paralogs in the pgPEN library. Red dashed line indicates the mean PC9 copy number of pgPEN library paralogs. Copy number data was obtained from DepMap.org. **(J)** Box and overlaid dot plot of CRISPR scores for sKO paralog target genes, grouped by PC9 copy number. Box plots indicate median ± IQR for each group. **(K)** Box and overlaid dot plot of CRISPR scores for dKO paralog target gene pairs, grouped by combined PC9 copy number. Box plots indicate median ± IQR for each group.

**Supplemental Figure 2.**
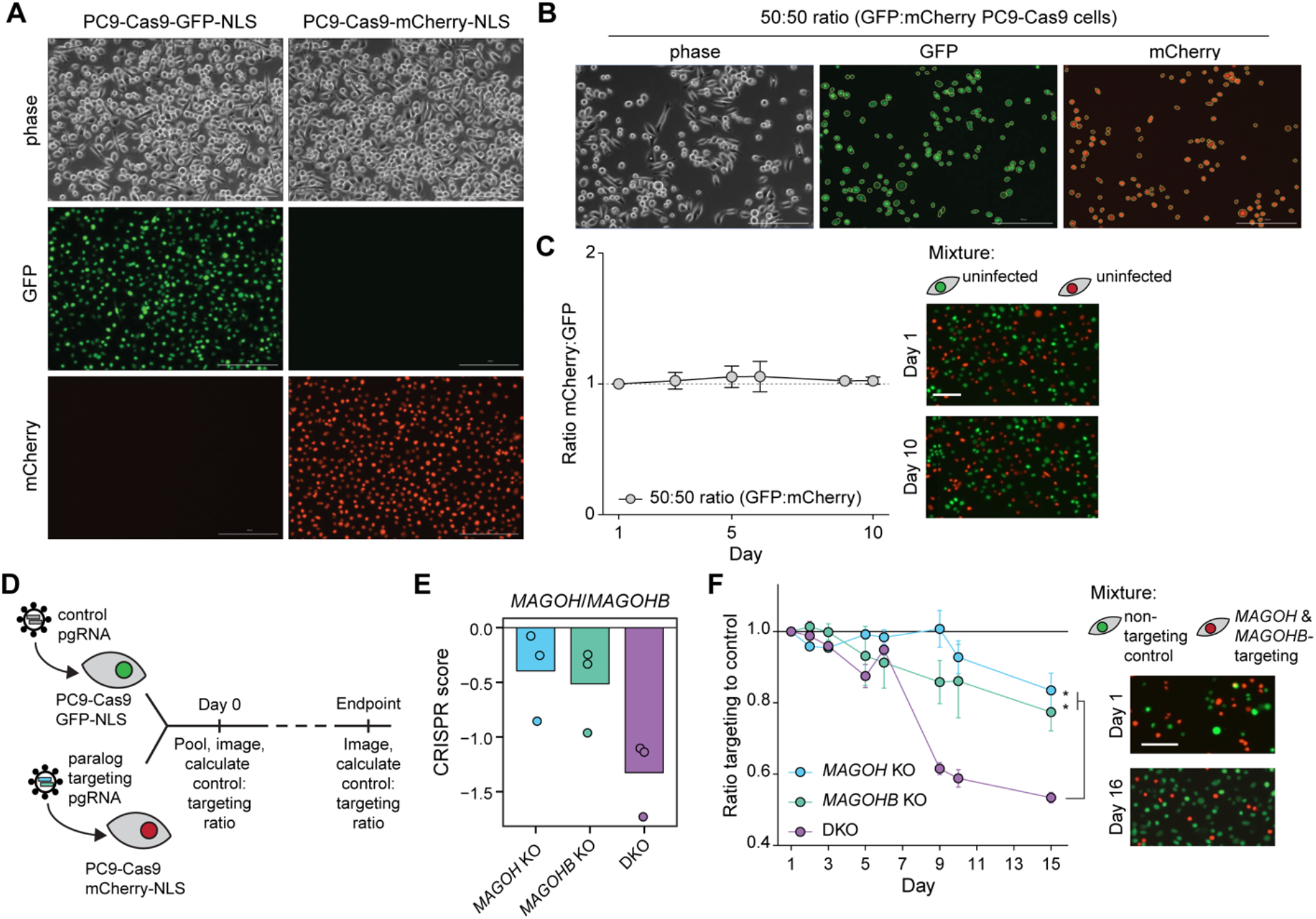
**(A)** Representative images of PC9-Cas9-GFP-NLS and PC9-Cas9-mCherry-NLS cells. **(B)** Representative images of detecting GFP and mCherry positive cells within a 50:50 mixture. **(C)** Line graph of PC9-Cas9-GFP-NLS and PC9-Cas9-mCherry-NLS cell relative viability across a 10 day time course. Representative images of mixture at day 1 and day 10 time points are shown. Scale bar, 100 μM. **(D)** Schematic of competitive fitness assay. **(E)** CRISPR scores for *MAGOH/MAGOHB* paralog pair in the PC9 screen. Data shown is the mean CS for each sKO or dKO target across three biological replicates with replicate data shown in overlaid points. **(F)** Line graph of relative viability for PC9-Cas9-mCherry-NLS cells expressing the indicated pgRNA compared to PC9-Cas9-GFP-NLS cells expressing a non-targeting control pgRNA. MAGOH/MAGOHB was selected as a positive control based on prior reports of synthetic lethality (Viswanathan et al., 2018). * *P* < 0.033 by one-tailed *t*-test. Right, representative images of mixture at day 1 and day 10 time points are shown. Scale bar, 100 μM.

**Supplemental Figure 3.**
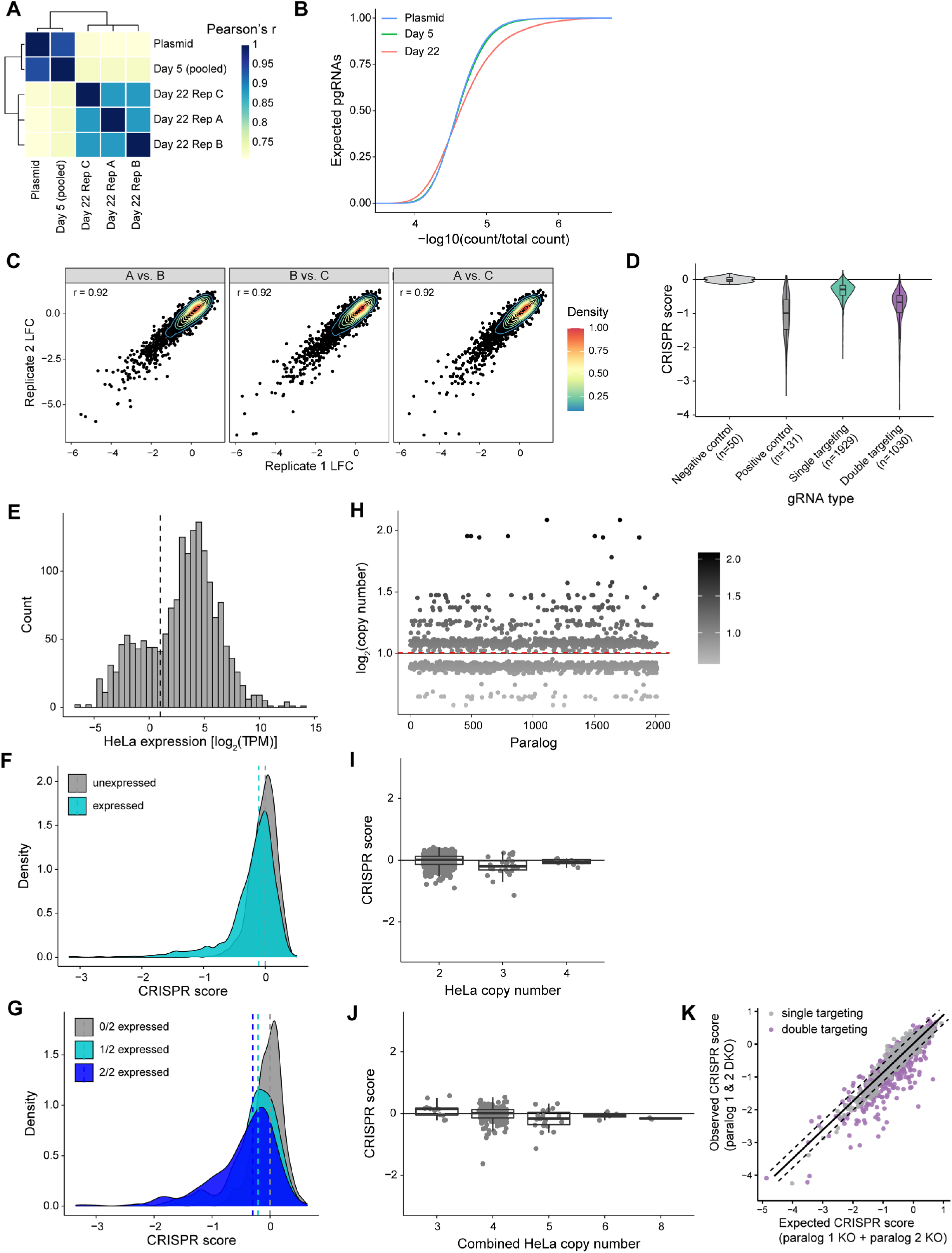
**(A)** Heat map illustrating Pearson correlations between sequencing reads in counts per million (CPM) supporting each pgRNA for all samples from the HeLa screen. Dendrogram, unsupervised clustering of CPM by the complete-linkage method. **(B)** Plot showing the cumulative distribution of pgRNA counts at each time point (mean across three biological replicates) in the HeLa screen. Each point represents the proportion of total pgRNAs with the corresponding count in a given sample. **(C)** Scatter plot of target-level log2(fold changes) between the plasmid and late time points for the indicated replicate comparisons in the HeLa screen. Contour lines indicate the density of points in 2D space. **(D)** Violin plots of target-level CRISPR scores for negative control (double non-targeting control), positive control (sKO pgRNAs targeting known essential genes), all other sKO, and dKO pgRNAs in the HeLa screen. **(E)** Histogram of PC9 RNA-seq reads in log2(TPM) for all paralogs in the pgPEN library. Dashed line indicates the cutoff used for expression: log2(TPM) > 1. **(F)** Density histogram of CRISPR scores for sKO paralog target genes, grouped by target gene expression in HeLa. **(G)** Density plot of target-level CRISPR scores for double targeting pgRNAs grouped by gene expression in HeLa cells indicating for each paralog pair whether zero, one, or both targeted genes are expressed (TPM > 2). Dashed lines indicate the median CRISPR score for each group. **(H)** Scatter plot of HeLa log2(copy number) for paralogs in the pgPEN library. Red dashed line indicates the mean HeLa copy number of pgPEN library paralogs. Copy number data was obtained from DepMap.org. **(I)** Box and overlaid dot plot of CRISPR scores for sKO paralog target genes, grouped by HeLa copy number. Box plots indicate median ±IQR for each group. **(J)** Box and overlaid dot plot of CRISPR scores for dKO paralog target gene pairs, grouped by combined HeLa copy number. Box plots indicate median ±IQR for each group. **(K)** Scatter plot of target-level observed versus expected CRISPR scores in the HeLa screen. Dashed line represents two residuals below the linear regression line for the negative control (single targeting) pgRNAs.

